# Recruitment and organization of ESCRT-0 and ubiquitinated cargo via condensation

**DOI:** 10.1101/2021.08.23.457352

**Authors:** Sudeep Banjade, Lu Zhu, Jeffrey R. Jorgensen, Sho W. Suzuki, Scott D. Emr

## Abstract

The general mechanisms by which ESCRTs are specifically recruited to various membranes, and how ESCRT subunits are spatially organized remain central questions in cell biology. At the endosome and lysosomes, ubiquitination of membrane proteins triggers ESCRT-mediated substrate recognition and degradation. Using the yeast lysosome/vacuole, we define the principles by which substrate engagement by ESCRTs occurs at this organelle. We find that multivalent interactions between ESCRT-0 and polyubiquitin is critical for substrate recognition at yeast vacuoles, with a lower-valency requirement for cargo engagement at endosomes. Direct recruitment of ESCRT-0 induces dynamic foci on the vacuole membrane, and forms fluid condensates *in vitro* with polyubiquitin. We propose that self-assembly of early ESCRTs induces condensation, an initial step in ESCRT-assembly/nucleation at membranes. This property can be tuned specifically at various organelles by modulating the number of binding interactions.

**One-Sentence Summary:** Condensation of multivalent ESCRT-0/polyubiquitin assemblies organizes cargo sorting reactions at lysosomes

## Main Text

ESCRT family members consist of a group of proteins involved in controlling diverse membrane-remodeling biochemical reactions, which include multivesicular body biogenesis, HIV budding, cytokinesis and membrane repair (*1*). Many of these occur at different membrane locations in the cell (endosome, lysosome, plasma membrane and the nucleus). Although the molecules that recruit ESCRTs are mostly known, the general principles of recruitment are less well understood. The mechanisms by which ESCRT reactions are spatially controlled remain unclear. In addition, while the structure of the individual subunits of the ESCRT complexes have been in most part solved (*2–5*), the mesoscale structural organization of early ESCRTs on the membrane is unclear.

At endosomes during multivesicular body biogenesis, ubiquitinated membrane proteins (cargo) recruit ESCRT-0, which then recruits the downstream proteins ESCRTs I, II and III(*6*). Therefore, ubiquitination serves as a signal to recruit ESCRTs to the endosomal membrane. The ESCRT-I complex has two ubiquitin binding motifs, ESCRT-II has one ubiquitin binding motif (*7*), and the complex of ESCRT-0 contains at least five ubiquitin binding motifs (one VHS and two UIM motifs in the ESCRT-0 protein Vps27, and one VHS and one UIM motif in the ESCRT-0 protein Hse1, (*7*)). Therefore, the ESCRT-0 complex is a critical component of the pathway that controls cargo binding. Furthermore, ESCRT-0 has been reported to form tetramers, oligomers and clusters (*8, 9*). *In vivo* cargo sorting occurs at hotspots (*10, 11*), demonstrating cargo concentration at specific sites, although the mechanism of this organization is not understood. The presence of multivalency for ubiquitin binding in ESCRT-0, and the knowledge of the existence of poly-ubiquitinated cargo as substrate, suggest higher order assembly of these complexes, as demonstrated for various multivalent interactions (*12–14*).

In this study, to understand the principles of ESCRT and cargo organization at membranes, we set out to delineate the properties of ESCRT’s initial recruitment and assembly at two similar but separate organelles in yeast endosomes and vacuoles (yeast lysosomes). We recently described the mechanism of ESCRT-mediated lysosomal protein recognition and degradation via ubiquitination of lysosomal membrane proteins (*15*). While studying the requirements of vacuolar membrane protein recognition by ESCRTs, we find that polyubiquitination is critical for efficient ESCRT recognition at the vacuole. Higher valency of ubiquitin is more critical for cargo sorting at the vacuole than at endosomes. Polyubiquitinated proteins associate with ESCRT-0, leading to self-assembly of ESCRT-0 and cargo into dynamic condensates, which may provide a platform for nucleation of downstream ESCRT complexes.

As a model cargo, we used the vacuolar protease carboxy-peptidase-S (CPS), which is transported to the vacuole from the Golgi via the endosomal pathway(*16*). GFP-CPS is sorted into multivesicular bodies by the action of ESCRTs and is then trafficked to the vacuolar lumen for degradation (Fig. 1A). Therefore, GFP-CPS is localized in the vacuolar lumen (Fig. 1D, Fig. S1A). In ESCRT mutants (for example in the ESCRT-0 mutants *vps27*Δ or *hse1*Δ), GFP-CPS gets stuck at aberrant endosomes (also called class-E compartments or “E-dots”) (Fig. S1 B).

**Fig. 1.**
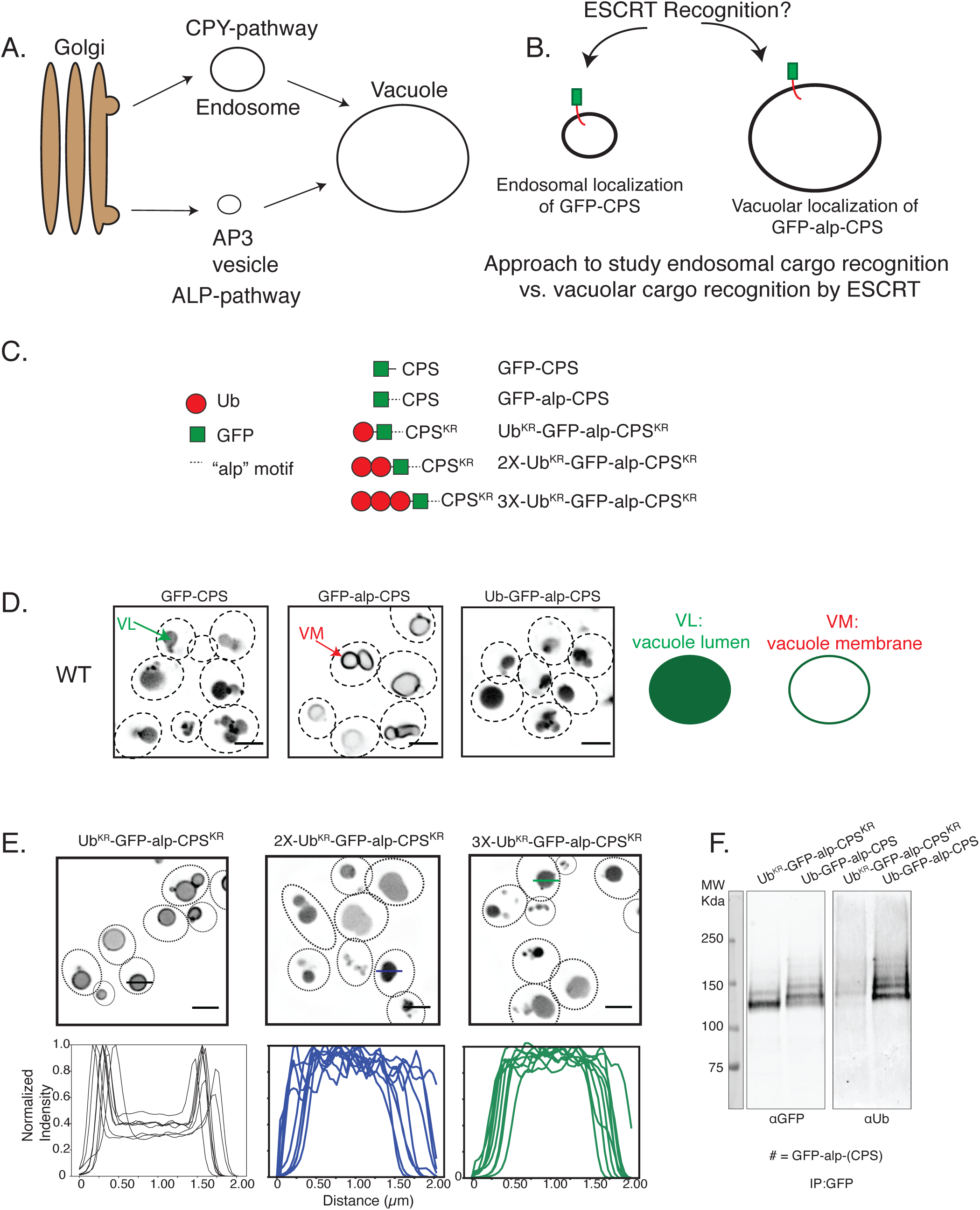
Polyubiquitination is required for efficient ESCRT-dependent cargo sorting at the vacuole. **A)** Model of carboxy peptidase S (CPS) and ALP trafficking to the vacuole. CPS traffics through the “CPY-pathway”/endosome where MVBs are formed by ESCRTs. The ALP-pathway forms separate vesicles that traffics proteins like the alkaline phosphatase to the vacuole. **B)** Model for the localization of GFP-CPS and GFP-alp-CPS at two sites where ESCRTs can function, providing a tool to study the same cargo at two membrane locations in the cell. **C)** Different CPS constructs used in this experiment. **D)**Localization of the CPS constructs as listed on top of the figures in a WT strain. The dotted lines represent outlines of yeast cells as imaged through the DIC channel. For simplicity the DIC channel is not shown but some images with the DIC channel are shown in the supplements (Fig S1). Note that VL denotes vacuolar lumen, VM denotes vacuole membrane. Right: A model of different variants of GFP-CPS localizing in the vacuolar lumen(VL) or vacuolar membrane (VM). Scale bars are 2 μm long. **E)** Localization of the GFP-CPS constructs as denoted in the figure. UB^KR^ has all the lysines mutated to Arg. Intensity profiles on the bottom represent line scan across the vacuoles. Representative lines are shown across three vacuoles (black, blue and green lines) in the microscopy images. Scale bars are 2 μm long. **F)** Immunoblots of UB^KR^-GFP-alp-CPS^KR^ (lysine-less substrate) and Ub-GFP-alp-CPS after performing immunoprecipitation with anti-GFP. These experiments were performed in a *doa4*Δ *pep4*Δ *prb1*Δ background strain.

While Golgi-endosome-vacuole is one route for proteins to get to the vacuole lumen, another route to the vacuole is through the AP-3 pathway (Fig. 1A). When GFP-CPS is re-routed to the vacuole membrane by adding an AP-3 recognition site (hereafter called GFP-alp-CPS – “alp” for the “alp pathway”), the protein remains at the vacuole membrane (Fig. 1C-D, (*16*)). These two proteins (GFP-CPS and GFP-alp-CPS) therefore provide us with almost identical proteins that get to the vacuole via two independent vesicular compartments. This design allows us to probe the properties that ESCRTs utilize for the same protein to be recognized at two different physical locations in the cell.

Our previous study indicated that ESCRTs are able to recognize ubiquitinated membrane proteins at the vacuole membrane as well, in addition to endosomes (*15*). Considering that GFP-alp-CPS is stable at the vacuole membrane and does not get internalized into the lumen, we tested if conjugating CPS with ubiquitin induces ESCRT-mediated cargo internalization. We found that when a single ubiquitin is conjugated at the N-terminus of GFP-alp-CPS, the protein now gets internalized into the lumen, in a Vps27-dependent fashion (Fig. 1C-D, Fig. S1B).

Following this observation, we asked if a single ubiquitin is sufficient for complete internalization of GFP-alp-CPS on the vacuole membrane. Therefore, we used a lysine-less ubiquitin molecule (where all the seven lysines are mutated to arginines) to probe the effect of a polyubiquitin-deficient cargo. CPS normally gets ubiquitinated on K8 and K12 residues (*17*), which were mutated to Arg in this construct, and from hereon called Ub^KR^-GFP-alp-CPS^KR^ (Fig. 1C).

We found that the internalization of this polyubiquitin-deficient cargo is inhibited and a large fraction of it remains at the vacuole membrane (Fig. 1E). Compared with the normal ubiquitin containing lysines, the lysine-less protein does not form polyubiquitin chains (Fig. 1F). If the number of the lysine-less Ub molecules is increased making 2X-Ub^KR^ or 3X-Ub^KR^, internalization to the lumen is rescued (Fig. 1C-1D).

Vacuole-targeted membrane-protein cargoes are ubiquitinated through the action of E3 ligases such as Rsp5, Tul1 and Pib1 (*18*). Deletions of Rsp5-adaptors Ssh4 and Ear1, in addition to Tul1 and Pib1, had a severe defect in internalization of the vacuolar-membrane cargo Ub-GFP-alp-CPS (Fig. S2). These data provide evidence that a higher level of ubiquitination (or polyubiquitination) is required for the vacuolar membrane protein to be recognized and sorted by ESCRTs.

The data argue that the vacuolar membrane proteome is susceptible to regulation through the ESCRT pathway and that polyubiquitination of cargo is necessary for efficient sorting and degradation. To study the effect of the number of ubiquitin molecules on vacuolar protein cargo in a controlled fashion, we utilized a previously established rapamycin-dependent degradation system (*15*). In this system, the protein/cargo of interest is tagged with FKBP, and the yeast strain also contains an FRB molecule conjugated with ubiquitin. This system allowed us to control the number of ubiquitin molecules conjugated to the cargo of interest by following the kinetics of cargo sorting and degradation after addition of rapamycin, which triggers FRB-FKBP binding.

A 2X-FKBP (two FKBP molecules fused in tandem) in the presence of FRB-3XUb (three ubiquitin molecules in tandem) was effective in inducing ESCRT-mediated sorting of the vacuolar lysine-transporter Ypq1 (*19*). We modified the valency of ubiquitin bound to the cargo by mutating one of the FRB-binding sites in FKBP (called FKBP*), and by using single or triple ubiquitin fused to FRB (Figure 2A). In cargo sorting assays, we found a strong dependence on the number (valency) of ubiquitin fused to FRB (Fig. 2A-D, Fig. S3A-C). The same effect is also observed when we used an orthogonal cargo Vph1, with the number of effective ubiquitin molecules controlling the level of internalization into the vacuolar lumen (Fig. S3C).

**Figure 2.**
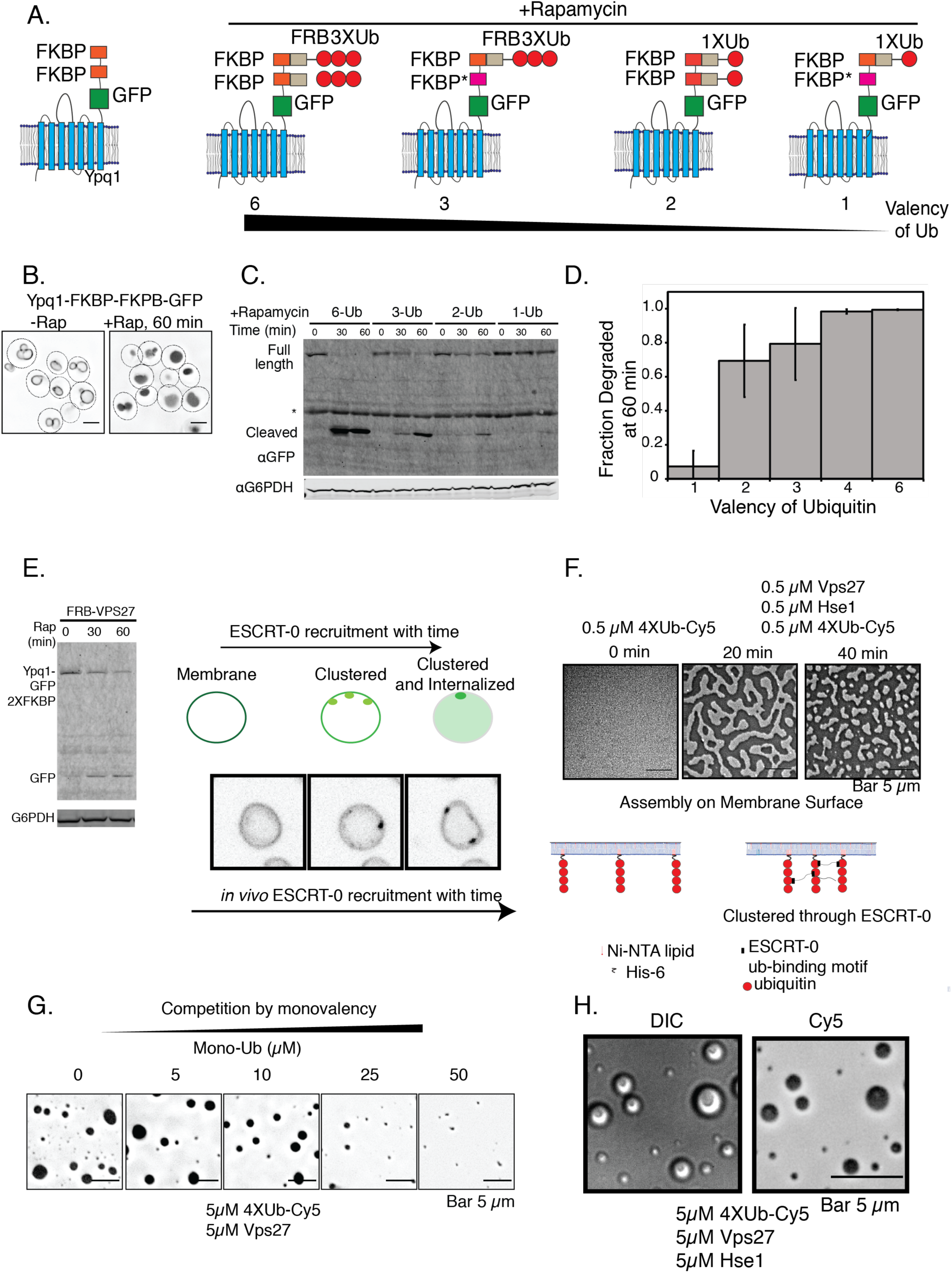
Multivalent interactions drive cargo clustering and sorting at the vacuole. **A)** Models of Ypq1 the FRB constructs used in B-D. **B)** Localization of Yqp1 constructs with and without rapamycin addition for 60 minutes. FKBP* contains mutations on the FRB-binding sites. **C)** Immunoblots for GFP after addition of Rapamycin for the indicated times. The free GFP band appears and increases in intensity as the full-length protein gets internalizes into the vacuole lumen. **D)** Quantification of the fraction GFP degraded into the vacuole lumen, at 60 minutes, from the data in **C). E)** Recruiting FRB-Vps27 onto the vacuolar cargo Ypq1-GFP-2XFKBP induces degradation of the cargo. **E)** Recruitment of FRB-Vps27 onto the vacuole via binding to Ypq1-GFP-2XFKBP also induces formation of foci on the surface of the vacuole. **F)** Assembly of Vps27-Hse1 and Cy5 labeled His6-4XUb on supported lipid bilayers consisting of 2% Ni-NTA PE. Bottom model-figure shows how 4X-Ub is associated with lipid bilayers. Model of ubiquitin binding motifs represent only a fraction of the ESCRT-0 complex. **G)** Competition experiment of the condensates with the inclusion of mono-ubiquitin at the mentioned concentrations. Protein components were combined together and fluorescence of Cy5 imaged after 2 hours at room temperature. Experiments were performed in 25 mM Bis-Tris pH 6.5, 150 mM NaCl. **H)** Recombinant Vps27 and Hse1 were incubated with 4X-Ub-Cy5 and imaged after 2 hours at room temperature, showing DIC image on the left and fluorescence (of Cy5) on the right. Experiments were performed in 25 mM Bis-Tris pH 6.5, 150 mM NaCl.

To assess whether direct recruitment of ESCRTs to the vacuolar cargo can cause degradation of the cargo, we fused the ESCRT-0 protein Vps27 to FRB. This construct is fully functional and identical to the activity of the wild-type Vps27 (Fig. S4), as it can fully complement the defect of a *vps27*Δ strain for two different cargo sorting reactions (Mup1-pHluorin and Can1 through a canavanine sensitivity assay, (*20*)).

Upon rapamycin-induced recruitment of Vps27 in this system, the modified cargo Ypq1 (fused to FKBP) internalized into the vacuolar lumen and degraded over time (Fig. 2E). Interestingly, we observed formation of distinct foci of the cargo at the vacuole surface upon adding rapamycin (Fig. 2E. Fig. S5). These foci are dynamic in nature – they move over time (Movie 1, Fig. S5), and occasionally fuse with one another. Some clusters disappear over time, as the cargo gets internalized into the lumen (Movie 1, Fig. S4C).

When a lower valency cargo construct is used instead, where one of the FKBP sites is mutated to abrogate FRB-Vps27 binding, upon rapamycin addition cargo clustering and sorting is inhibited (Fig. S6A). Importantly, this lower-valency cargo molecule is able to bind to FRB-Vps27 in a rapamycin-dependent fashion (Fig. S6B). Therefore, while ESCRT-0 is recruited, the valency of binding sites on the Ypq1 construct is critical to induce of ESCRT-mediated cargo sorting.

Our data therefore suggest that multivalent interactions between ESCRT-0 and cargo induce clustering, sorting and degradation. To understand the basis of cargo clustering induced by ESCRT-0, we purified the ESCRT-0 proteins Vps27 and Hse1, and reconstituted the assembly reaction *in-vitro*, using His6-tagged ubiquitin as the model cargo. We fused four ubiquitin genes in tandem with one another and labeled this molecule with Cy5. Upon assembling this model ubiquitin with ESCRT-0, micron sized condensates formed on supported lipid bilayers and in solution at low micromolar concentrations (Fig. 2F-H). The concentrations required for formation of condensates is lower in the nanomolar range on supported lipid bilayers and micromolar range in solution (Fig. 2F), implying robust formation of these assemblies. The structures also exhibit morphologies expected of liquid-like structures on membranes (Fig. 2F,(*12*)).

Inclusion of a single ubiquitin in the assay abrogated condensate formation, signifying the importance of ubiquitin valency in the self-assembly (Fig. 2G). A wide range of condensate sizes, down to nanometer scale are formed, as depicted by electron microscopy (3A). Rough edges of the condensates down to the nanometer level, and highly dense meshwork of proteins are reminiscent of network formation in such assemblies, as previously observed in other multivalent systems (*14, 21*).

These structures are dynamic since they exhibited molecular exchange ((Fig. 3B), fluorescence recovery after photobleaching). Solvent conditions are important for their formation, as condensation was favored by acidic pH (Fig. S7A,). These properties are hallmarks of biomolecular condensates, structures that exhibit dynamic rearrangement of molecules and rapid molecular exchange with the environment (*13*). These condensates did not form with only 4XUb, or with 4XUb and Hse1 (Fig. S7B). Without Hse1, the condensates of 4X Ub and Vps27 were smaller (Fig. S7B), as the valency of interaction is reduced. Condensate formation was possible but significantly inhibited with only Vps27 and Hse1, signifying weak self-association between the subunits of the ESCRT-0 complex (Fig. S7B), probably owing to the higher-order oligomerization and intrinsic disorder of the complex (Fig. S6C).

**Figure 3.**
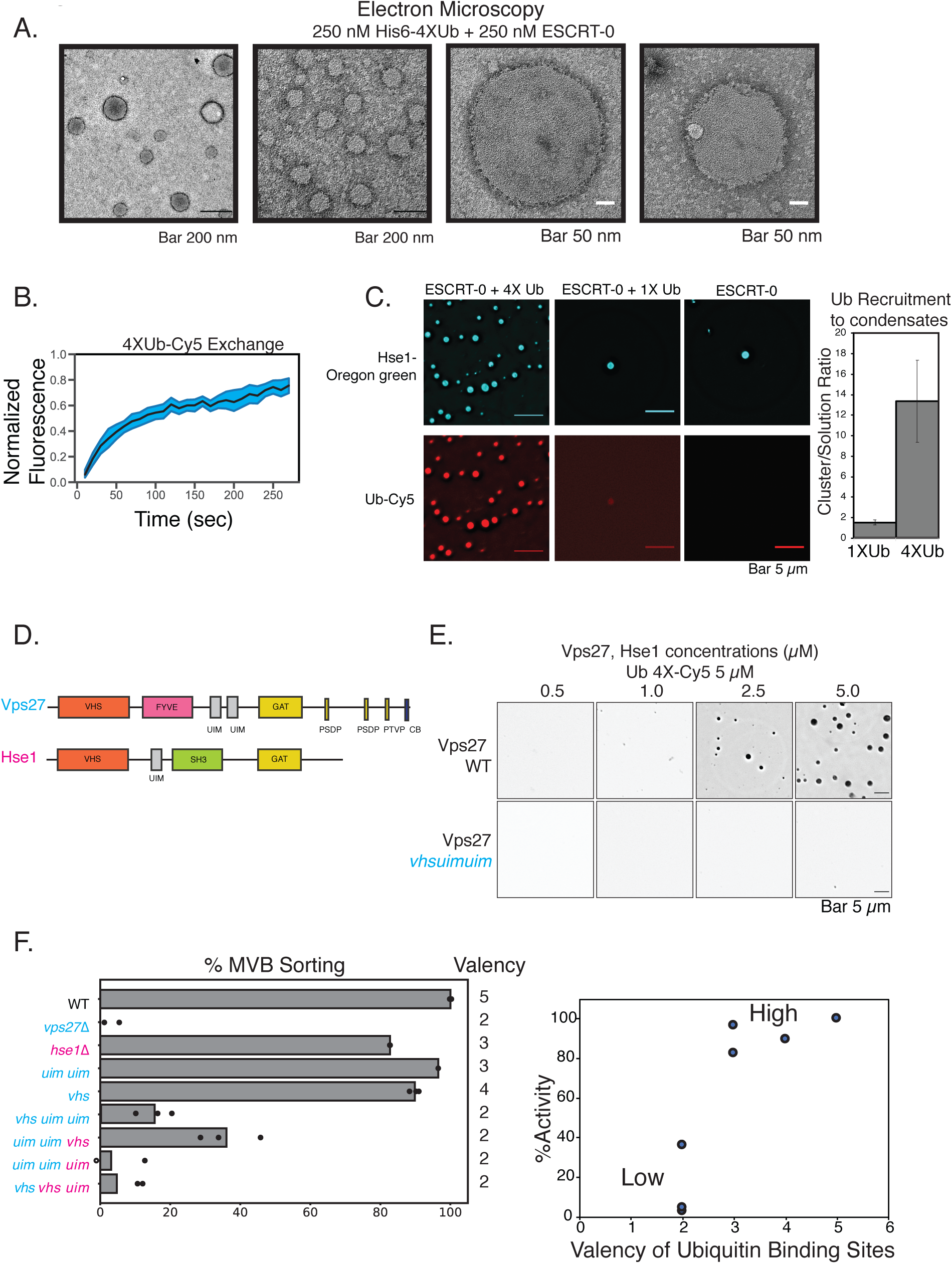
Multivalent interactions drive cargo sorting reactions. **A)** Electron microscopy images of the ESCRT-0 complex (Vps27/Hse1) with His6-4X-Ub. The structures form variable sizes distributions and are spherical in nature. Electron microscopy was performed at 25 mM Hepes pH 7.5, 150 mM NaCl. **B)** Fluorescence recovery after photobleaching (FRAP) of 4XUb-Cy5. Three whole droplets were bleached to obtain the normalized fluorescence recovery curves. **C)** Purified Vps27 and Hse1 were incubated with 4X-Ub and imaged after 2 hours at room temperature. 4X-Ub contains a His6 tag at the N-terminus and also a cysteine that has been labeled with a Cy5 dye for visualization. Hse1 was labeled with Oregon-green maleimide. Right figure is a quantification of the level of mono-ubiquitin-Cy5 or tetraubiquitin-Cy5 recruitment in droplets made from respective proteins. **D)** Domain organization of the ESCRT-0 proteins Vps27 and Hse1. Vps27 contains three ubiquitin binding domains (a VHS and two UIMs), while Hse1 contains two (a VHS and a UIM motif). **E)** Condensate formation with WT Vps27 or the Vps27 *vhsuimuim* mutant, in the presence of Hse1 and 4XUb-Cy5. Experiments were performed in 25 mM Bis-tris pH6.5, 150 mM NaCl. The mutant Vps27 contains mutations in the VHS domain and mutations in the UIM motifs that abrogate ubiquitin binding. **F)** Mup1-pHluorin sorting assay through flow cytometry with different ESCRT-0 mutants. Mutations were made in the ESCRT-0 components Vps27 and Hse1 at their VHS and UIM motifs. Cyan italics represent mutations in Vps27, and the magenta italics represent mutations in Hse1. Numbers on the right of the panel represent the number of ubiquitin binding sites present in the ESCRT-0 complex in the various mutants. Dots in the bars represent independent experiments, while the bars represent average sorting from three independent experiments. Right: Sorting activity (as obtained from F) for Mup1-phluorin sorting as a function of ubiquitin valency in ESCRT-0.

Mutations in the ubiquitin binding motifs have valency dependent effects on cargo sorting assays with the model endocytosis cargo Mup1 (Fig. 3D, 3F). Mutations in the ubiquitin-binding motifs of Vps27 (Vps27^*vhs uim uim*^) also inhibited condensate formation *in-vitro* (Fig. 3E).

Our data therefore imply that the role of multivalent interactions during cargo sorting is to cluster cargo into condensates. Higher valency interactions through polyubiquitin and ESCRT-0 is also critical for cargo sorting reactions at the vacuolar membrane for several cargoes (GFP-alp-CPS, Ypq1, Vph1) and also for endosomal cargo (Mup1). Interestingly, the internalization of the lysine-less GFP-CPS molecule, (Ub^KR^-GFP-CPS^KR^: note without the alp signal), that traffics through the endosome, localizes to the lumen of the vacuole (Fig. 4A), in contrast to the vacuolar cargo (Fig. 1). Therefore, the cargo recognition and internalization of this polyubiquitin-deficient membrane protein is more severely defective at the vacuolar membrane compared to the endosomal membrane. These effects are observed also when we follow the kinetics of degradation of the vacuole-targeted cargo compared with that of the endosome-targeted cargo. We used a galactose inducible expression system, following the stability of the protein over time upon translation inhibition with cycloheximide (Figure 4A). Although the proteins reach the endosome/vacuole membrane (Figure 4C), the kinetics of degradation of the vacuole targeted protein is much slower than that of the endosome-targeted protein (Fig. 4A-B). The rate of degradation can be enhanced with a 3XUb^KR^ conjugated molecule (Fig. S8). These data imply that lower valency interactions are more efficient for cargo sorting at endosomes than at vacuolar membrane.

**Figure 4.**
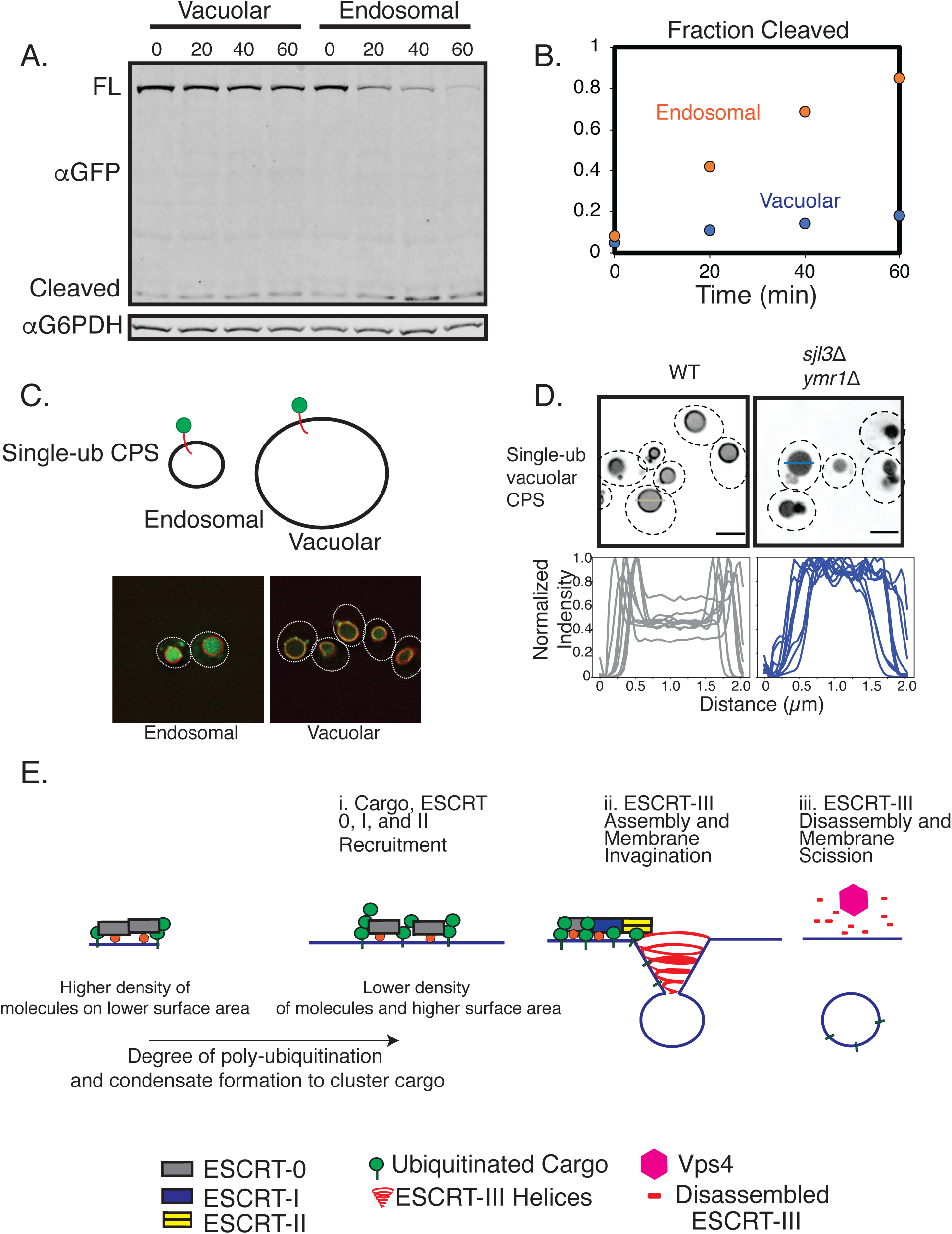
Differential dependence of valency on cargo sorting on endosomal and vacuolar cargo. **A)** Kinetics of the degradation of UB^KR^-GFP-CPS^KR^ (endosomal) and UB^KR^-GFP-alp-CPS^KR^ (vacuolar) proteins upon induction of expression for 20 minutes and further inhibition of translation with cycloheximide addition. The 0 timepoint represents 20 minutes of GAL induction with no cyloheximide. **B)** Quantification of the fraction of GFP cleaved of the data from **A). C)** Localization of the UB^KR^-GFP-CPS^KR^ (endosomal) and the UB^KR^-GFP-alp-CPS^KR^ (vacuolar) constructs after inducing the expression for 20 minutes through the GAL promoter and then stopping translation through cycloheximide addition for 30 minutes. **D)** Localization of the indicated UB^KR^-GFP-alp-CPS^KR^ (vacuolar cargo) in the phosphatase mutant strains in the phosphatidyl-inositol-3P phosphatase mutant *ymr1*Δ *sjl3*Δ. Intensity profiles on the bottom represent line scan across the vacuoles. Representative lines are shown across two vacuoles (gray and blue lines) in the microscopy images. Scale bars are 2 μm long. **E)** Models suggesting how variation of ESCRT and cargo density affects recruitment and condensation of early ESCRT and substrate on the surface of membranes. The condensation of early players in the pathway may nucleate downstream assembly, leading to ESCRT-III polymerization and membrane budding.

Endosomes are sites of high density of ESCRTs (*22*). We reasoned that due to the higher density of ESCRTs at the endosomes, there is a lower requirement for multivalent interactions to recruit and condense ESCRTs at this organelle, as opposed to the vacuolar membrane. One of the lipid-species that recruit ESCRTs to the endosomal membrane is phosphatidyl-inositol-3P (PI3P). We hypothesized that an increase in PI3P density could in turn increase the density of ESCRTs, thereby reducing the requirement of higher valency interactions at the vacuolar membrane. To test this prediction, we used the PI3P phosphatase mutants *ymr1*Δ and *sjl3*Δ, which have previously been shown to increase the level of PI3P on the vacuole membrane (*23*). In the single mutants ymr1Δ and *sjl3*Δ, the single-ubiquitin conjugated vacuolar GFP-CPS (Ub^KR^-GFP-alp-CPS^KR^) remains mostly on the vacuole membrane (Fig. S9B). However, in the double mutant *ymr1*Δ *sjl3*Δ, the molecule is enriched in the vacuole lumen (Fig. 4D). Therefore, the defective sorting of the substrate can be rescued by the local increase in density of PI3P at the vacuolar membrane.

Our data provide evidence for two distinct mechanisms of recruitment and organization of ESCRT complexes at two separate membrane locations: the endosomes and the vacuole (Fig. 4D). The general principles behind function of ESCRTs at membranes were proposed to include - recruitment to membranes, “self-assembly”, and ESCRT-III polymerization (*24*). Our data suggest that one of the functions of the ESCRT-0 complex, by virtue of having multiple binding sites for ubiquitin, is to facilitate cargo clustering at the membrane through biomolecular condensation. ESCRT-0 by itself has a weaker ability to self-associate and undergo condensation. When substrates are polyubiquitinated, the multivalency of ESCRT-0 enhances formation of higher-order species of the ESCRT-0/cargo complexes. Increasing the density of PI3P increases the local concentration of ESCRT-0/ubiquitin on the membrane, thereby lowering the threshold for self-association (Fig. 4D, S10).

Condensing cargo via early ESCRTs could provide multiple advantages in the cargo-sorting process. The physical concentration of cargo could amplify sorting of cargo, increasing the specificity for cargo recognition. Concentrating downstream ESCRT complexes at a particular location and increasing the dwell time on the membrane could also provide a platform for nucleation of ESCRT-III polymerization, as observed for actin assembly pathways through upstream condensing nucleators (*25*). The rapid dynamics of the ESCRT-ubiquitin complexes also could allow for facile dissociation of the ESCRTs from cargo. Furthermore, condensation of ubiquitinated cargo could feed-back to the ubiquitin-ligase machinery, allowing for enhanced ubiquitination of the cargo, further providing specificity for cargo recognition.

At locations in the cell where ESCRT-0 is not involved, there may exist other mechanisms of self-assembly – in the case of HIV budding, the HIV Gag protein is known to form higher-order assemblies, which recruit downstream ESCRTs (*26*). Clustering and condensation of ESCRT-recruiters, therefore, could be a general property of ESCRT-related systems.

Our data provide clues for how clustering of signaling molecules can be regulated at various locations in the cell. By changing the number of binding sites for a complex on the membrane, cells may be able to quickly adjust local concentrations of enzymes and signaling adaptors. 2D surfaces and their charge properties may also play an important role in the nucleation of condensates (*27*) and also promote membrane repair and remodeling (*28, 29*). The composition of the membrane therefore should play a critical role in controlling formation and function of membrane-associated biomolecular condensates.

## Acknowledgments

We thank all the past and current members of the Emr lab for valuable discussions. Sudeep Banjade was an HHMI fellow of the Damon Runyon Cancer Research Foundation. This work was also supported by a Cornell University Grant to Scott D Emr.

## Author Contributions

Conceptualization: SB

Methodology: SB, LZ, JJ, SS

Investigation: SB

Funding acquisition: SB, SDE

Supervision: SDE

Writing – original draft: SB

Writing – review & editing: SB, SS, SDE

## Data and materials availability

All data are available in the main text or the supplementary materials. Strains and plasmids can be shared upon request.

## Supplementary Materials

Materials and Methods

Figs. S1 to S10

Table S1

Movie 1

**Movie 1: *In vivo* visualization of Ypq1-GFP-2X-FKBP upon addition of rapamycin**. Yeast cells were treated with 1 μg/mL of Rapamycin and imaged over the timeframe indicated in the movie.

## Supplementary Materials

### Materials and Methods

**Strains and Plasmids** are listed in Table 1.

**Table 1:**
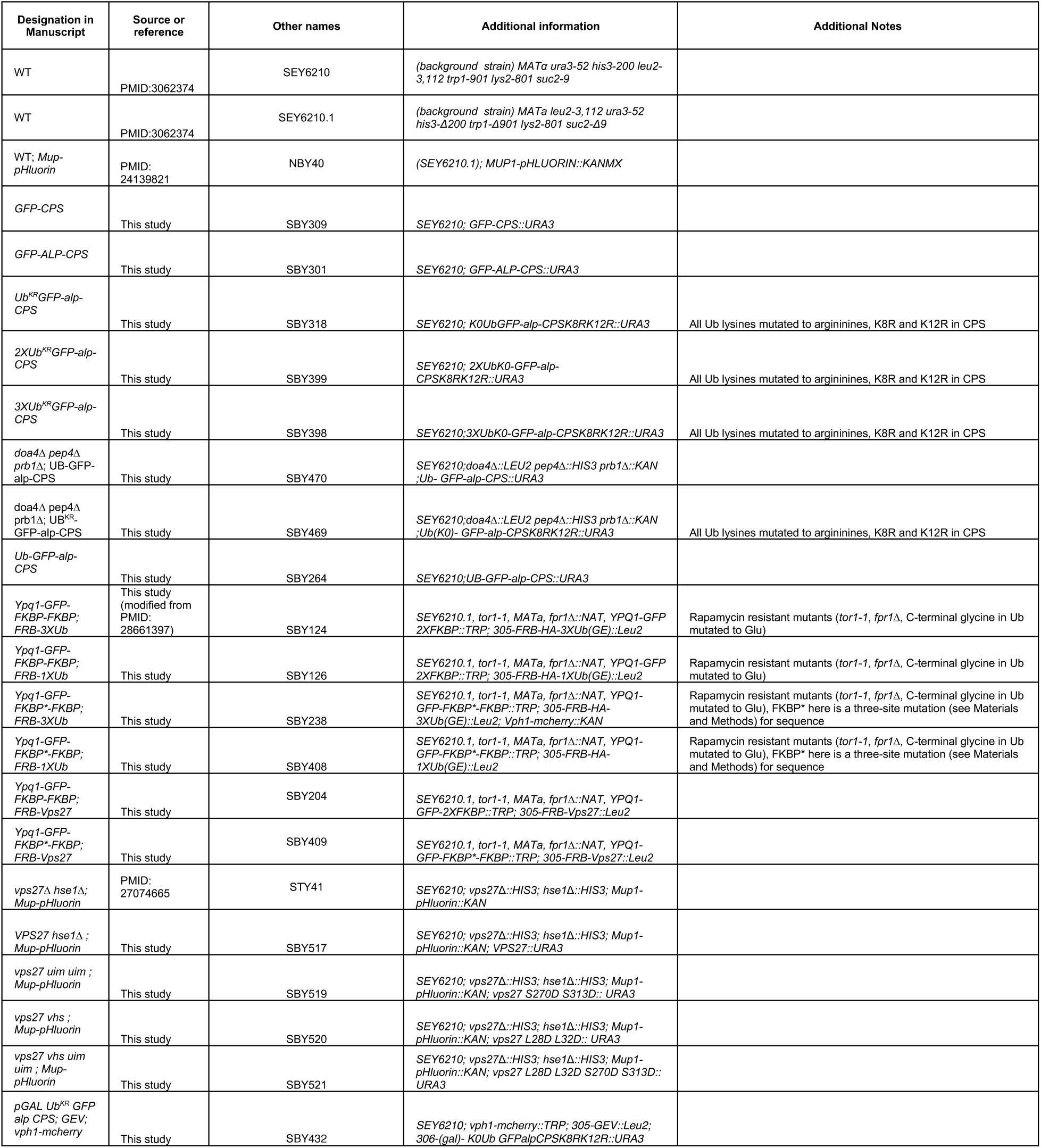
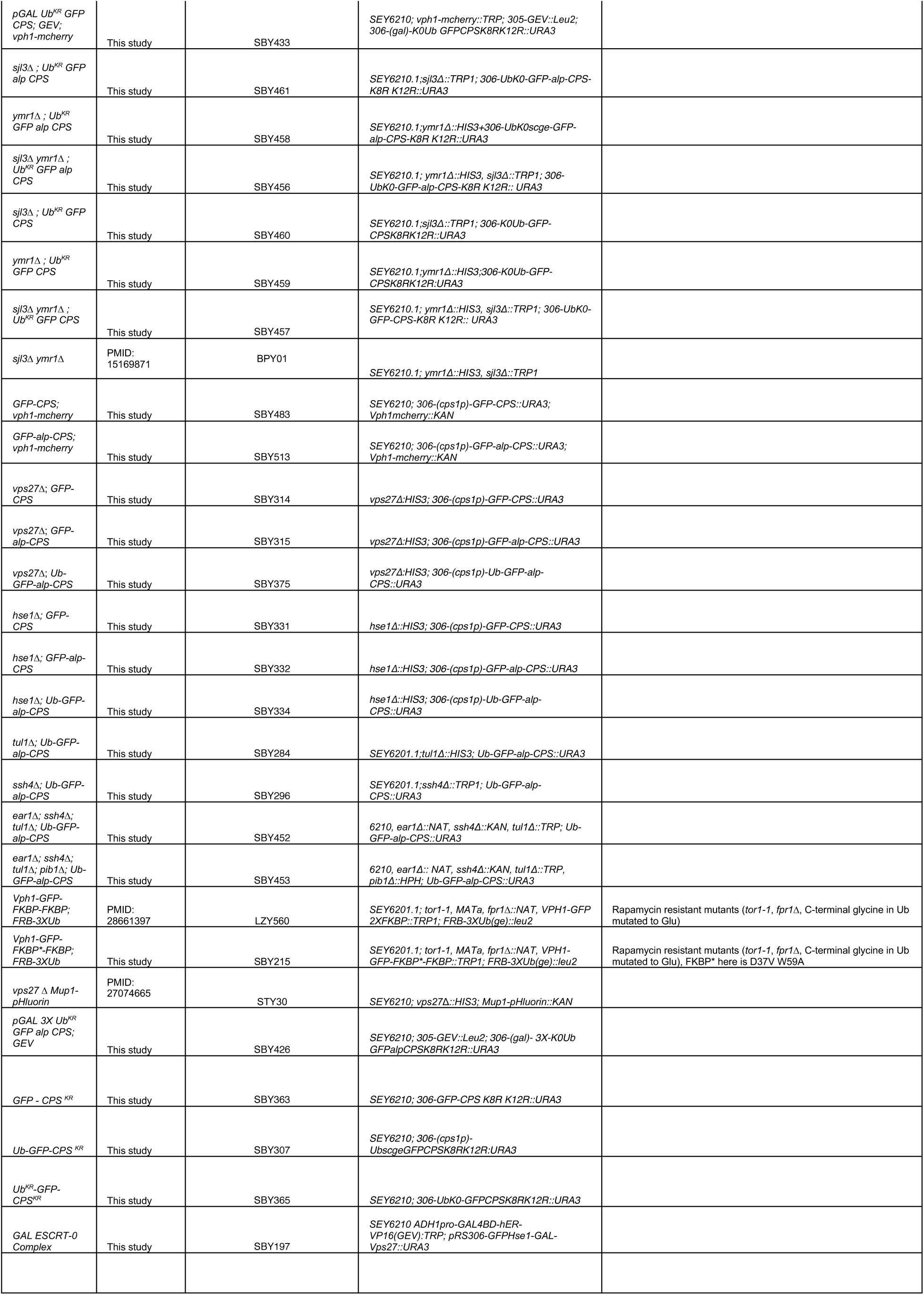
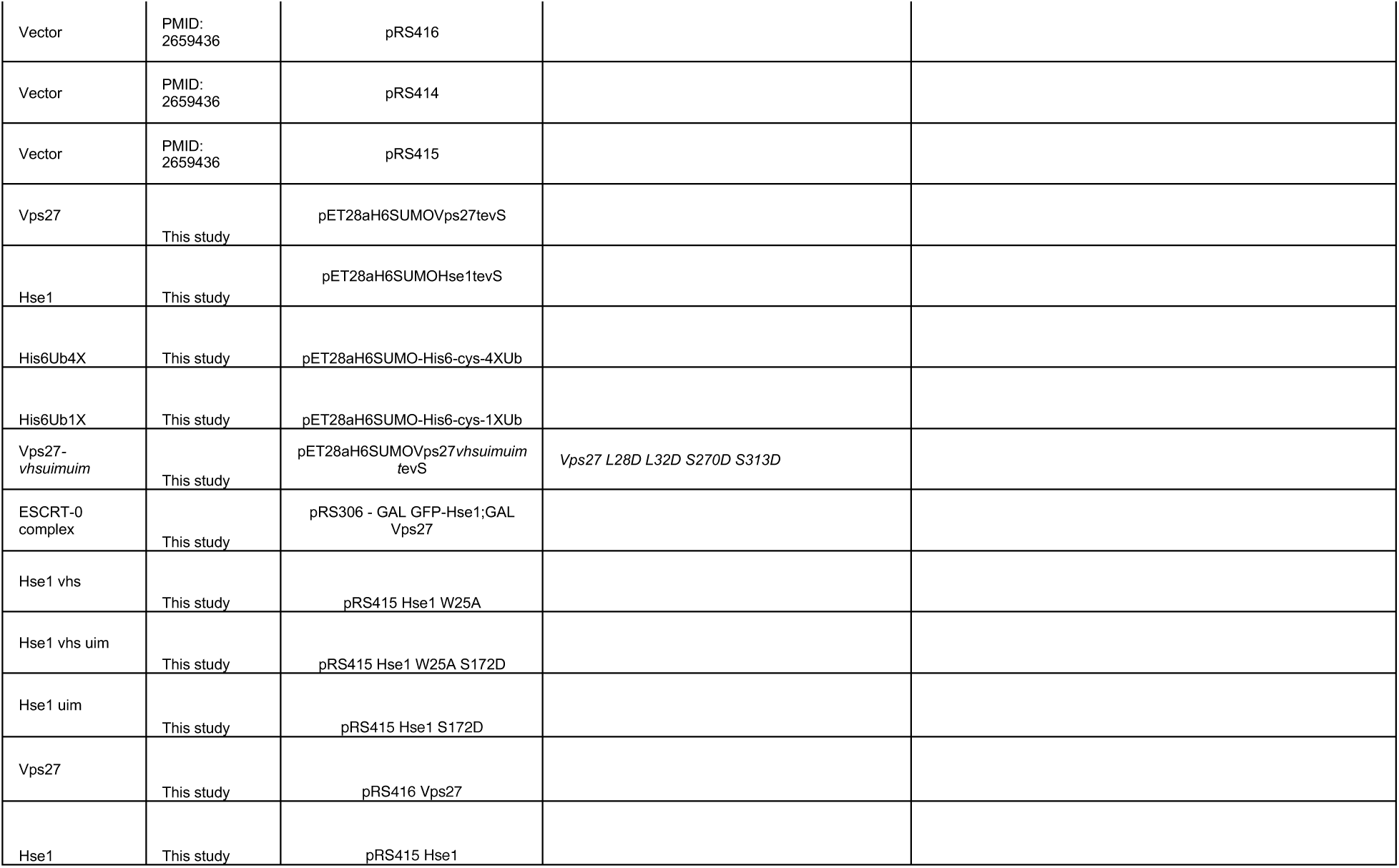
Strains and Plasmids.

### Fluorescence Microscopy

For microscopy purposes of steady-state distribution of GFP-CPS (and its derivatives), Ypq1-2XFKBP-GFP and Vph1-2XFKBP-GFP, the following procedure was used. Cells were grown on YPD (yeast extract, peptdone and dextrose mixture). 1 mL of mid-log cells were centrifuged for 1-2 min at 10,000 xg and resuspended in 10-20 μL of Milli-Q water. Imaging was performed on a Deltavision Elite system with an Olympus IX-71 inverted microscope using a 100X/1.4 NA oil objective. On the Softworx software, images were deconvolved using 10 iterations with the “conservative” method. Images were analyzed on Image J. Linescan images were analyzed after background subtraction and normalized to the range of 0 to 1 by dividing the intensities by the maximum intensity.

Time-lapse microscopy was performed as follows. Chambered slides were first rinsed with Milli-Q water. 200 μL of 1 mg/mL concanavalin A was applied to the glass slide for 1 minute at room-temperature. Excess concanavalin A was removed and the slide rinsed twice with 1 mL Milli-Q water. Mid-log cells were centrifuged for 1-2 min at 10,000 xg and resuspended with minimal medium. Cells were incubated on the slides for 20 minutes, then rinsed twice with 1 mL of minimal medium. Imaging was then performed on a CSU-X spinning-disk confocal microscopy system (Intelligent Imaging Innovations), coupled to a DMI 6000B microscope (Leica), 100X/1.45-NA objective and a QuantEMCCM camera. Analysis was performed on Slidebook and Image J.

### GAL Induction and Cycloheximide Chase Assay

GFP-CPS constructs with a GAL promoter consist of an estradiol-responsive element (*30*). For expression of the protein, cells were grown to mid-log, then 100 nM of β-estradiol was added to the medium for GAL induction. After 20 minutes, translation was stopped by adding 50 μg/mL of cycloheximide. Samples were taken (5 ODs of cells) 20 minutes after adding β-estradiol, and then at subsequent times as indicated in the figures.

### Rapamycin mediated FKBP-FRB interactions

Different versions of FRB-Ub, FRB-Vps27 and membrane proteins (Ypq1, Vph1) were allowed to interact through rapamycin-triggered FKBP-FRB association. All rapamycin treatment assays were done similarly by adding 1 μg/mL of rapamycin to mid-log cells for different amounts of time before imaging or taking samples for immunoblots. FKBP* was made by mutating three different Rapamycin-FRB binding sites to ensure complete abrogation of FRB binding in our multivalent system. The sequences of human FKBP and FKBP* are provided below with the mutations highlighted in bold letters:

FKBP:

MGVQVETISPGDGRTFPKRGQTCVVHYTGMLEDGKKFDSSRDRNKPFKFMLGKQEVIRGWEEGVAQMSVGQRAKLTISPDYAYGATGHPGIIPPHATLVFDVELLKLE

FKBP*:

MGVQVETISPGDGRTFPKRGQTCVVHYTGMLEDGKK**AVS**SRDANAPFKFMLGKQ**AAA** R**GA**EEGVAQMSVGQRAKLTISPDYAYGAT**AAAAA**IPPHATLVFDVELLKLE

### Immunoblots

Western blots were performed as follows, as described before (*31*). 5 OD equivalent of cells were collected by centrifugation at 4000 xg. After washing with 1 mL of cold H2O, and then centrifuged again at 4000 xg, cells were precipitated with 10% TCA for >1 hr on ice. Cells were washed twice with 1 mL of acetone, resuspending pellets between washes by bath sonication. Pelleted cells were then lysed in 100 μL lysis buffer (50 mM Tris-HCl, pH 7.5, 8 M urea, 2% SDS, and 1 mM EDTA) by bead beating for 10 min. 100 μL of sample buffer (150 mM Tris-Cl, pH 6.8, 8 M urea, 10% SDS, 24% glycerol, 10% v/v βME, and bromophenol blue) was then added to the sample and vortexed for 10 min. After centrifugation for 6 min at 21,000 xg, supernatant was loaded on an SDS-PAGE gel and transferred onto a nitrocellulose membrane. Imaging of the western blots was performed using an Odyssey CLx imaging system and analyzed using the Image Studio Lite 4.0.21 software (LI-COR Biosciences). Fractional degradation of the GFP tagged proteins were quantified by dividing the intensity of the GFP band from the sum of the intensities of the full-length and the cleaved-GFP bands.

### Denaturation IP for Ubiquitination Assay

Ubiquitination of the CPS constructs were performed in a strain background of *doa4*Δ *pep4*Δ *prb1*Δ. CPS constructs were integrated into this strain and ubiquitin was overexpressed using a myc-Ub construct under the control of a copper promoter. The copper promoter was induced by adding 100 μM CuSO4 for 4 hours to mid-log cells. 100 OD equivalent of cells were harvested and washed with Milli-Q water. Cells were resuspended in urea containing buffer (50 mM Tris-HCl pH 8.0, 1% SDS, 8 M urea, 20 mM NEM, 1X Roche protease inhibitor without EDTA). Cells were lysed with zirconia beads from Biospec (about 500 □L of bead volume), by vortexing twice for 30 s at 4° C. Cells were then chilled on ice for 10 minutes. An equivalent volume of a high-salt containing buffer (50 mM Tris-HCl pH 8.0, 500 mM NaCl, 10% glycerol, 20 mM NEM, 0.2% Triton X 100) was added to the lysate and chilled again on ice for 10 min. After centrifugation for 10 min at 16,00 xg, supernatant was incubated with anti-GFP beads (at a volume ratio of 1:50 beads:solution). This mixture was rotated for 2 hours at 4° C. Beads were washed with wash buffer (50 mM Tris-HCl pH 8.0, 250 mM NaCl, 0.5% SDS, 4 M urea, 20 mM NEM, 5% glycerol), 3 times equaling an 8000-fold dilution. Elution was performed by boiling the beads at 98° C for 3 minutes in 100 μL of sample buffer (150 mM Tris-Cl, pH 6.8, 8 M urea, 10% SDS, 24% glycerol, 10% v/v βME, and bromophenol blue). Samples were then blotted for GFP and ubiquitin.

### Canavanine Assay and Mup1-Sorting Assay

These assays were performed exactly as described before (*20, 31, 32*). Briefy, canavanine spot plates were performed on synthetic media with canavanine at various concentrations and imaged after 3-5 days. Mup1 sorting assay was performed on Mup1-pHluorin integrated cells after addition of 20 μg/mL at various time points. Fluorescence measurements were done on a BD Accuri C6 flow cytometer.

### Formation of Lipid Monolayers, Liposomes and Supported Lipid Bilayers

Monolayers were formed as previously described(*20, 31, 32*). Supported lipid bilayers were formed as follows, following previous protocols (*5*).

To form the supported bilayers, liposomes were first created by extrusion. Lipid mixtures of 2% Ni^2+^-NTA DOGS and 98% POPC were created in chloroform. These lipid mixtures were allowed to evaporate overnight in a desiccator. The dried mixture was hydrated in 25 mM Hepes pH 7.5, 150 mM NaCl for ∼ half an hour at room temperature with vortexing every 5 min. These multilamellar vesicles were used to create small unilamellar vesicles (SUVs) by extrusion through a 100 nm extrusion filter (Avanti).

To make supported lipid bilayers, 96-well plate glass bottomed plates (Cellvis) were used. Plates were cleaned by treating with 5% Hellmanex solution and incubating at 42 ° C for 30 minutes. After thorough cleaning with Milli Q water, the plates were then treated with 6 M NaOH at room-temperature for 2 hours. The plates were again thoroughly washed with Milli Q water. To make supported bilayers, the liposome mixture was warmed to 37 ° C and then applied to the cleaned glass-bottomed wells. After incubating at room-temperature for 10 minutes, unabsorbed vesicles were washed with BSA buffer (25 mM Hepes pH 7.5, 150 mM NaCl, 0.1% BSA). The plates were allowed to sit in this buffer for 30 minutes. Proteins were then added to this buffer and incubated for various times as indicated in the text for analysis of assembly on the membrane surface. Imaging was performed with either Deltavision widefield or a Leica Confocal microscope.

### Protein Purification

All proteins were expressed in the Rosetta strain. Lysis was performed by sonication on ice. Lysed cells were cleared by centrifugation at 26,000 x g for 40 minutes at 4 °C. Affinity purification of his-tagged proteins were performed using the Talon cobalt beads .

#### Ubiquitin

Ubiquitin constructs had an N-terminal His6-SUMO tag followed by a His6 tag and an N-terminal cysteine for maleimide-dye conjugation. 1X ubiquitin was expressed with 0.25 mM IPTG at 37 °C for 4 hours, and 4X Ub was expressed with 0.5 mM IPTG at 18 °C overnight. After affinity purification with cobalt beads, and cleavage of His6-SUMO on beads, 4X-Ub was passed through a Hi-Trap S-Sepharose cation exchange column, and the flow-through collected. 1X-Ub was purified with cobalt beads and H6-Sumo cleaved on beads. Subsequently, these ubiquitin proteins were treated with 1 mM fresh DTT, and then ran though SD200 in buffer lacking reducing agent for labeling. Proteins were concentrated and incubated with Cy3 or Cy5-maleimide (Lumiprobe) at 5-fold excess and room-temperature to label the proteins. Excess dye was removed by running through a Hi-Trap Q column and dialysis. Unlabeled proteins were similarly flash-frozen right after the SD200 step.

#### ESCRT-0

Vps27 and Hse1 constructs for expression were made in the pET28a vector containing an N-terminal His6 and SUMO tags and a C-terminal S-tag. Proteins were expressed at 18°C overnight with 1 mM IPTG. After Co^2+^ column, the SUMO tag was cleaved on beads by the protease ULP overnight at 4 °C. The cleaved protein was then applied to an S-column. The flow-through from this step was then concentrated and ran through an SD200 column. A substantial fraction of the Vps27 protein comes out in the void volume, which is not used in the study as the nature of this fraction of protein is not clear, but it is possible that this population also reflects a fraction that is self-assembling. We collect the fraction that enters the column for our studies. Hse1-oregon green was made using the same labeling protocol as ubiquitin. In this case, the endogeneous cysteines on Hse1 were used for labeling purposes. Our experiments with Hse1-oregon green include only 10% of the labeled protein.

Initial experiments with ESCRT-0 complex was also purified from *S. cerevisiae* using GAL promoters. In this approach, Hse1 was tagged with GFP at the N-terminus, followed by a Prescission cleavage site. GFP-prescission-Hse1 under a GAL promoter was expressed together with Vps27 (also under GAL promoter) in the same vector. This plasmid was integrated into a strain consisting of a Gal4-ER-VP16 system, whose expression can be induced by applying β-estradiol. This strain was grown at 30 deg C until mid-log, and induced with 100 nM β-estradiol until saturation. Cells were collected and lysed using a freezer mill. GFP-prescission-Hse1 was affinity-purified using a home-made GFP-nanobody agarose column. The Hse1-Vps27 complex was cleaved off by using Prescission protease on beads at 4 deg C for ∼16 hours. Cleaved protein was concentrated and purified further using an SD200 column (GE), in 25 mM Hepes pH 7.5, 150 mM NaCl and 10% glycerol. Protein was flash-frozen in liquid nitrogen and stored at -80 deg C. This complex was used for the electron microscopy experiments.

### Electron Microscopy

On the monolayer-grids, proteins were incubated for 1 hour. Grids were stained with 2% ammonium molybdate. Electron microscopy of the monolayers associating proteins was performed on an FEI Morgagni 268 TEM.

**Fig. S1.**
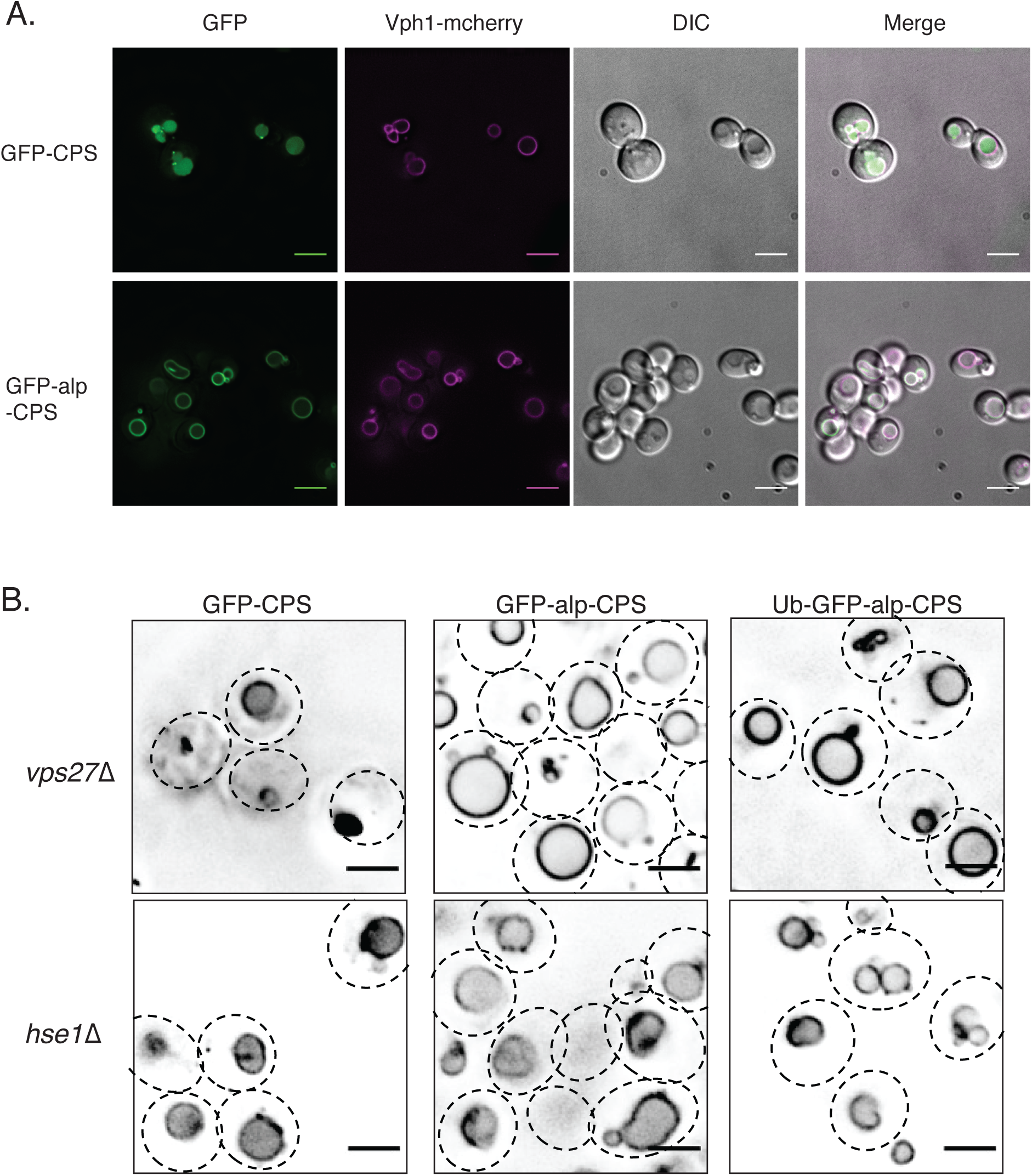
Localization of endosomal and vacuolar GFP-CPS molecules. A) GFP-CPS or GFP-alp-CPS in wild-type strains expressing Vph1-mcherry. Vph1 is a vacuolar membrane protein. Scale bars are 5 μm. B) Localization of different GFP-CPS constructs in the ESCRT-0 components *vps27*Δ and *hse1*Δ strains. Scale bars are 2 μm each.

**Fig. S2.**
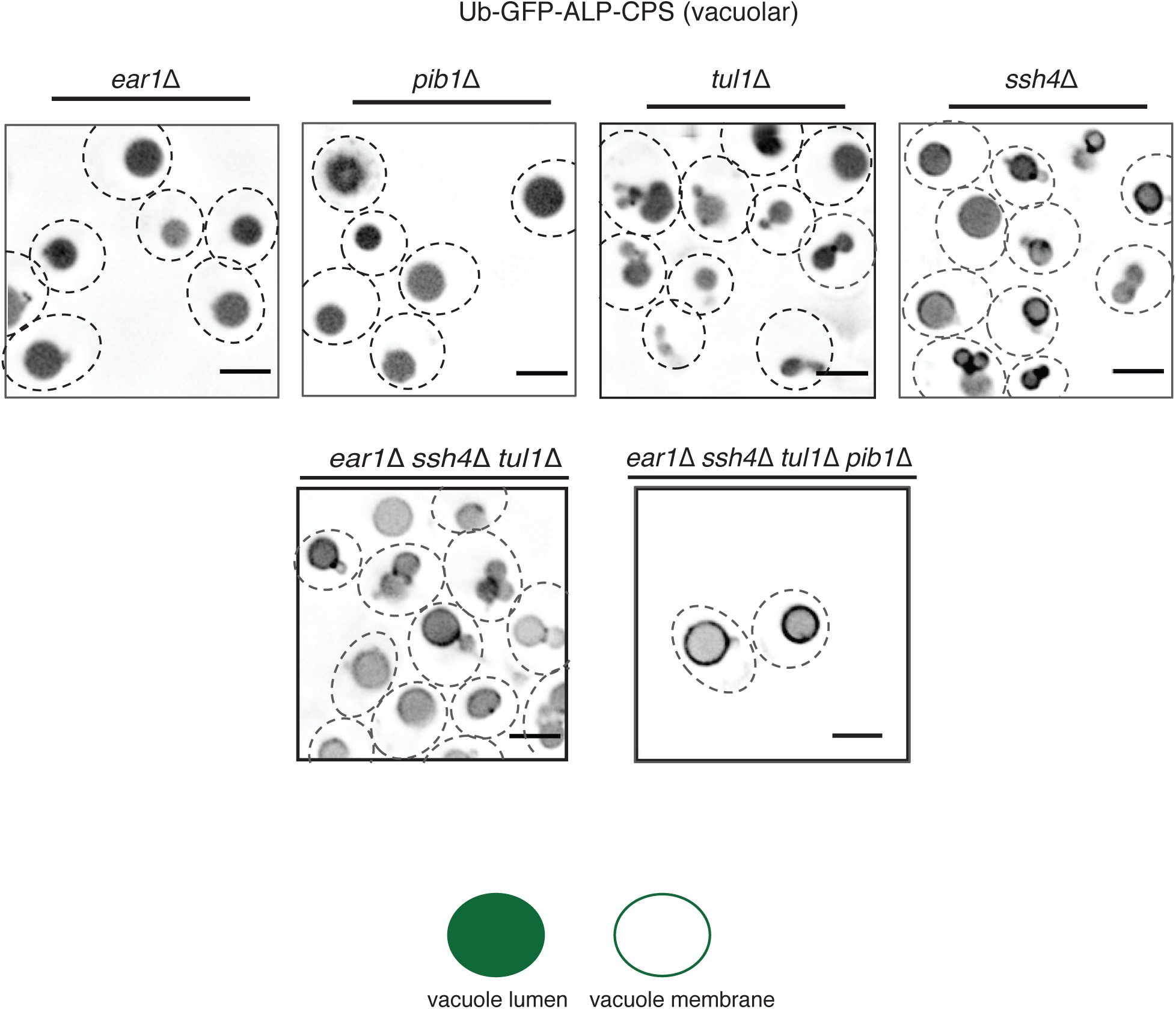
Poly-ubiquitination is critical for vacuolar sorting of ESCRT cargo. Localization of Ub-GFP-alp-CPS (ubiquitin conjugated vacuolar cargo) in the *ear1*Δ *ssh4*Δ *pib1*Δ *tul1*Δ or the individual strains. Scale bars are 2 μm each.

**Fig. S3.**
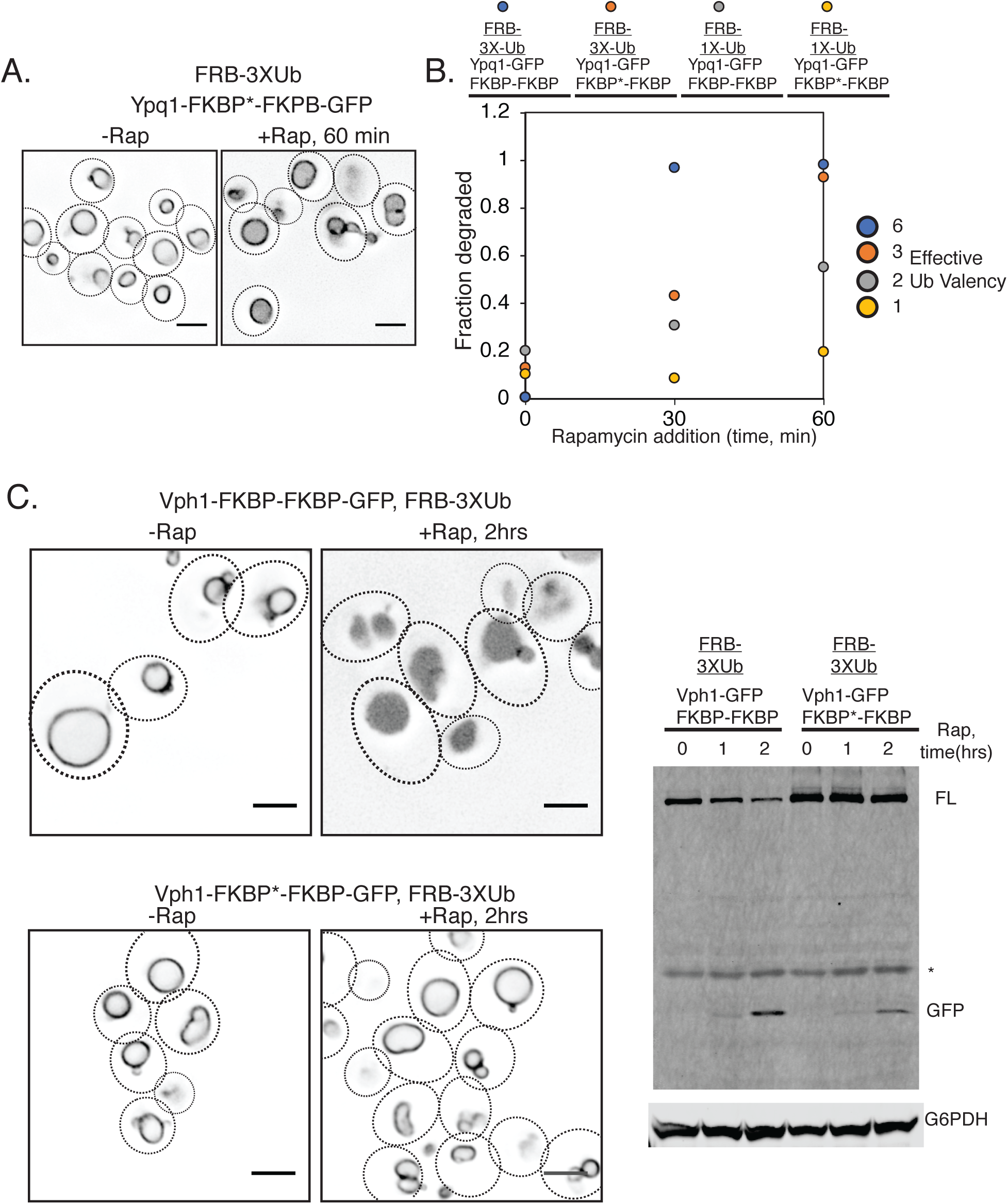
Multivalent interactions of polyubiquitin and ESCRT-0 promotes cargo sorting. **A)** Effect of rapamycin-induced sorting of Ypq1 construct with one FKBP site mutated and can no longer interact with FRB. Strain consists of FRB-3X ubiquitin and the imaging was done before adding rapamycin and after 60 minutes of adding rapamycin. **B)** Kinetics of different Ypq1-GFP-FKBP molecules in the presence of different FRB-Ub molecules. Colors of circles indicate the effective valency of ubiquitin achieved from the combination of FKBP and FRB constructs. **C)** Vph1-GFP-FKBP-FKBP constructs sorting in the presence of FRB-3XUb. Mutation in one of the FRB binding sites in FKBP inhibits sorting of the Vph1 construct, signifying the effect of multivalent interactions in sorting efficiency. Left figures are imaging experiments following GFP-tagged Vph1, and the right image is an immunoblot following GFP cleavage after adding rapamycin for the indicated times.

**Fig. S4.**
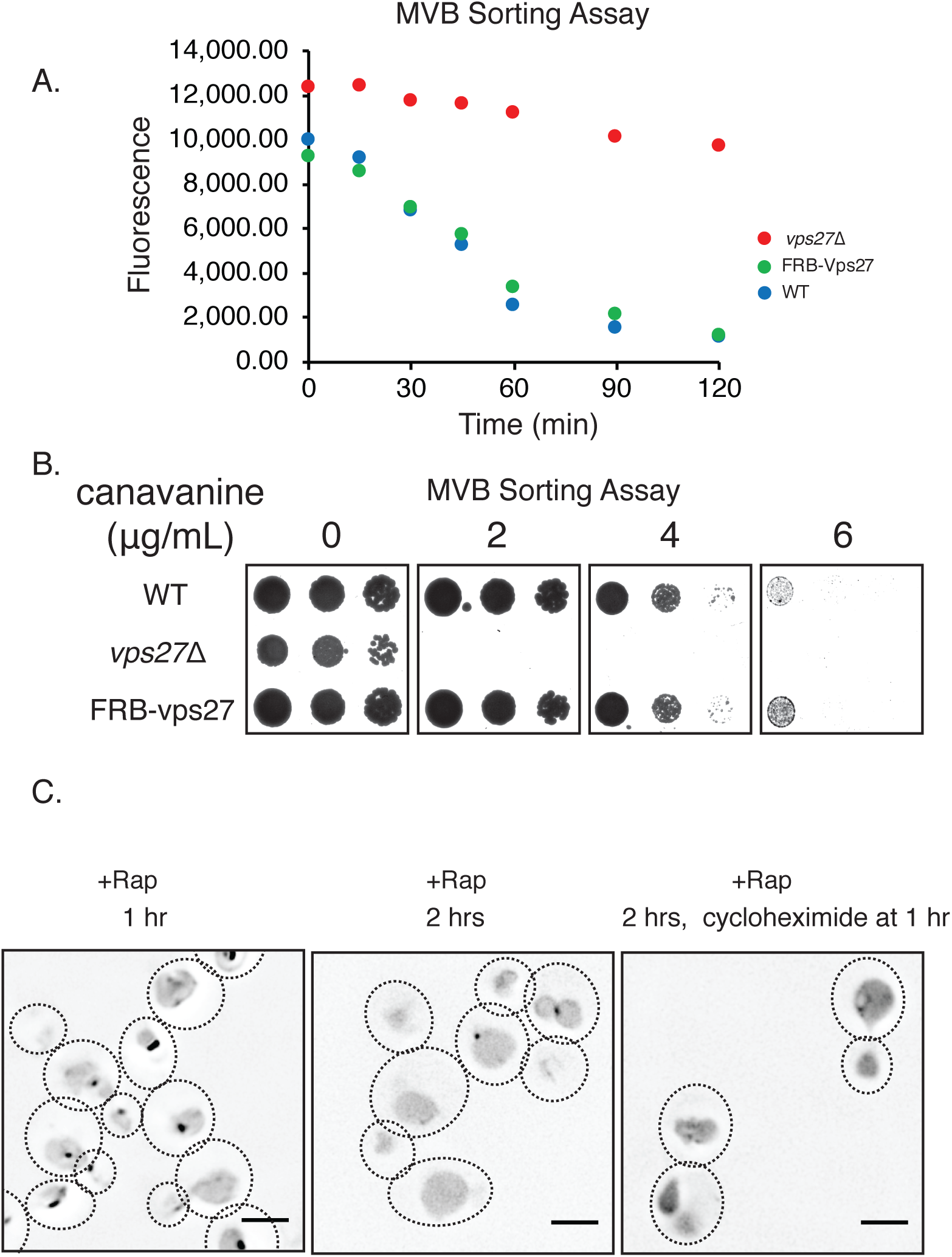
Recruitment of ESCRT-0 to vacuolar membrane induces cargo clustering. **A-B)** FRB-Vps27 construct used is identical to WT Vps27 in cargo sorting assays for the cargoes Mup1 in Mup1-pHluorin sorting **(A)** and Can1 sorting canavanine assays **(B). C)** With time or with translation inhibition, the clusters induced by FRB-Vps27 recruitment disappear, presumably by internalization into the vacuolar lumen.

**Fig. S5.**
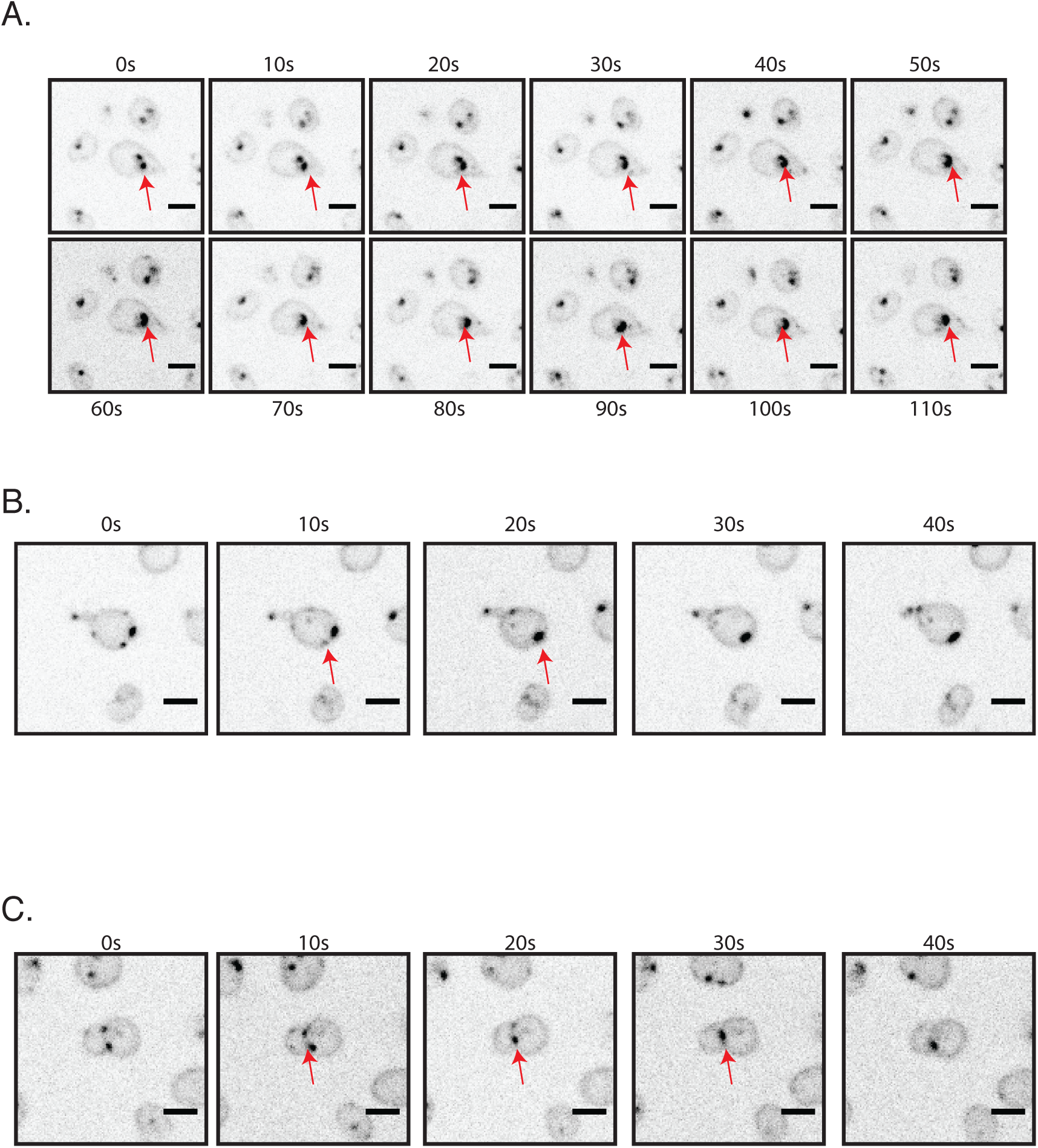
*In vivo* dynamics of clusters of Ypq1-GFP-2XFKBP molecule induced by recruitment of FRB-Vps27 (A-C). In these experiments, rapamycin was added for 30 minutes the imaging was performed with 10 ms interval. Scale bars are 2 μm each.

**Fig. S6.**
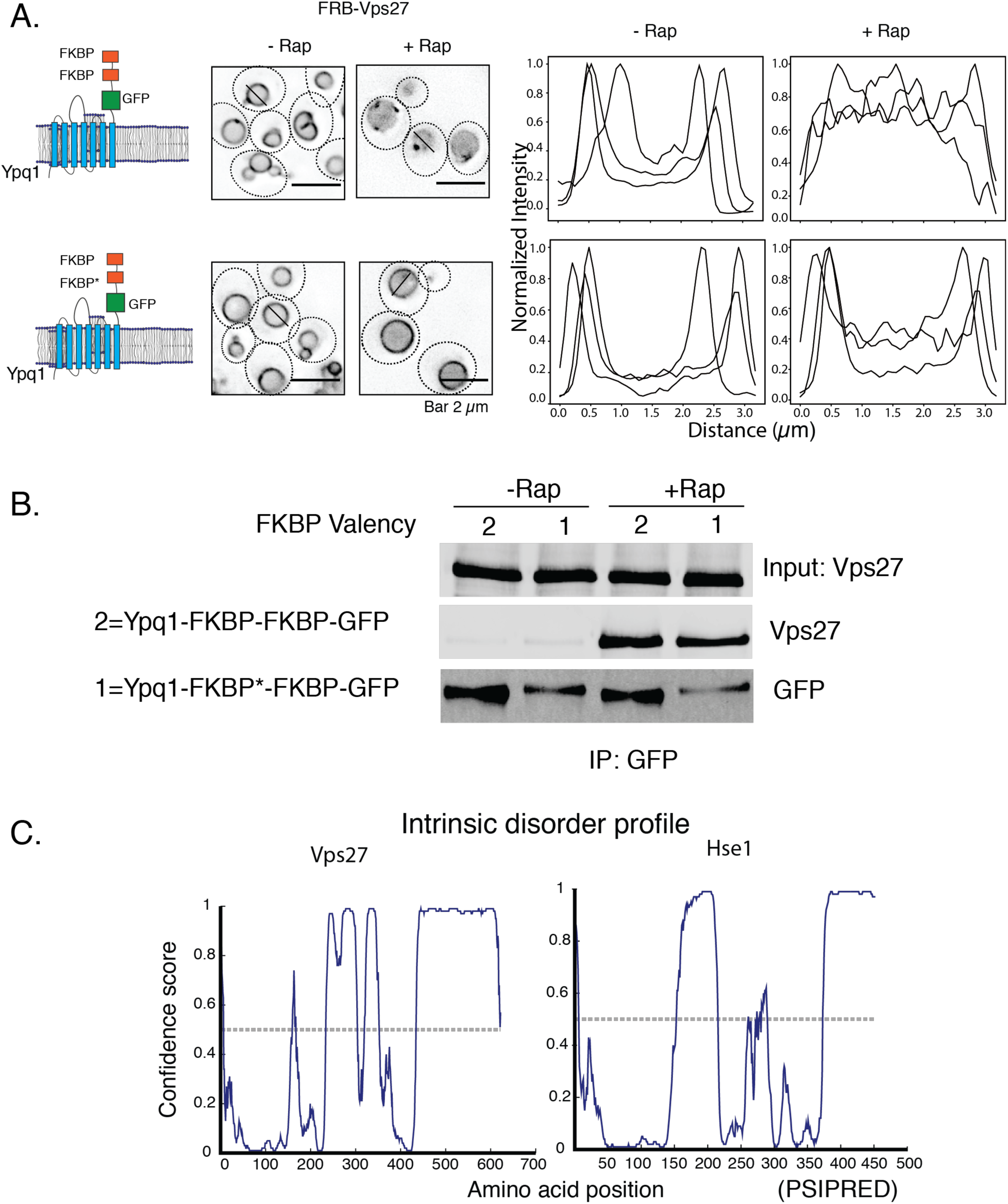
Valency effect on ESCRT-0 recruitment and cargo sorting. **A)** Ypq1-GFP-FKBP-FKBP or one of the FKBPs mutated to abrogate FRB binding were imaged after adding rapamycin to strains consisting of FRB-Vps27. Cargo clustering and internalization are reduced in the strain expressing the lower valency molecule. Graphs on the right are representative normalized intensity profiles of line-scans across vacuoles from microscopy images. **B)** Immunoprecipitation of GFP in Ypq1-GFP-FKBP-FKBP and FRB-Vps27 expressing strains and blotting for Vps27. FRB-Vps27 is still bound to Yqp1-GFP-FKBP*-FKBP in a rapamycin dependent fashion. **C)** Intrinsic-disorder profile obtained from DISOPRED for the ESCRT-0 components Vps27 and Hse1.

**Fig. S7.**
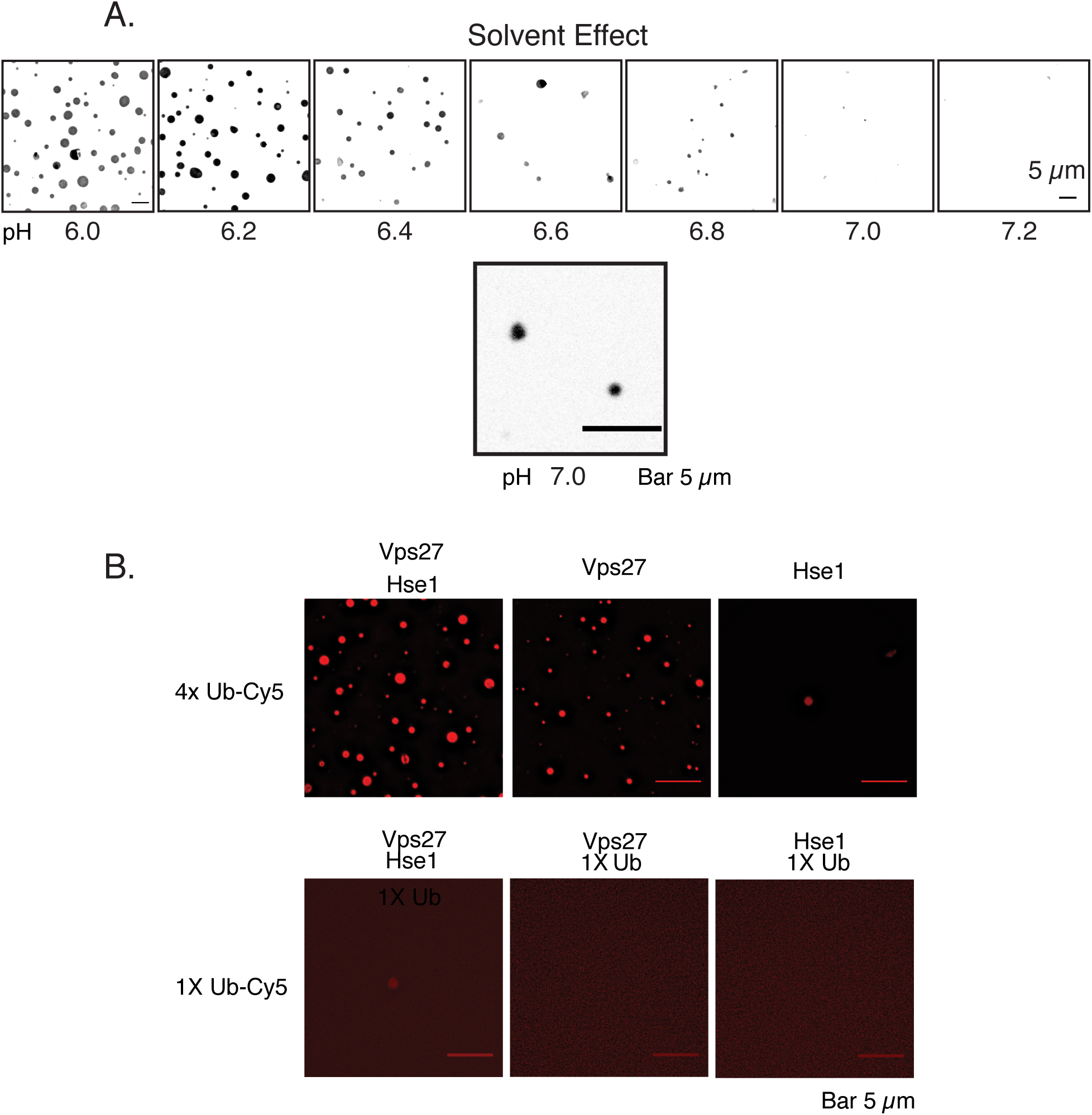
Condensation of ESCRT-0 and polyubiquitin assemblies. **A)** Solvent effect on the condensation of ESCRT-0 and 4X ubiquitin. pH was varied by using different buffers of Bis-Tris at the indicated pH values. Concentration of buffer was kept at 100 mM and NaCl was kept constant at 150 mM. Images were taken after two hours of assembling the components at room temperature. Bottom figure shows a zoomed-in image of the droplets at pH 7, showing presence of droplets but at reduced numbers. Scale bars are 5 μm each. **B)** Assemblies of ESCRT-0 and either 4XUb or 1X Ub at the indicated compositions (either both ESCRT-0 components or singles, as represented on the top of the panels). Concentrations of the molecules were at 5 μM, and the buffer was 25 mM Bis Tris pH 6.5, 150 mM NaCl.

**Fig. S8.**
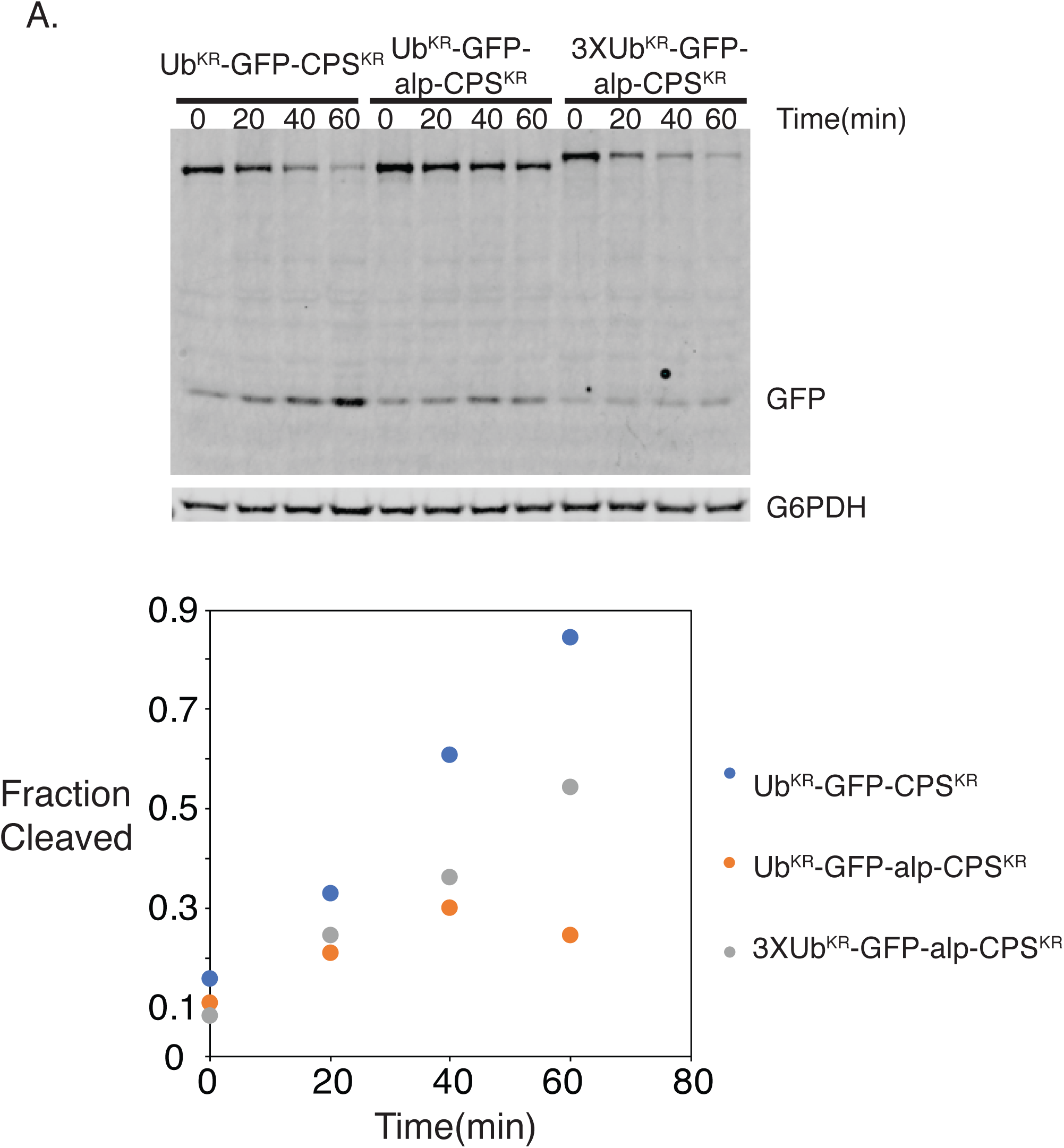
Enhancing ubiquitin chain length induces vacuolar cargo sorting. **A)** Kinetics of the degradation of Ub^KR^-GFP-CPS (endosomal), Ub^KR^-GFP-alp-CPS (vacuolar), or 3XUb^KR^-GFP-alp-CPS proteins upon induction of expression for 20 minutes and further inhibition of translation with cycloheximide addition. The 0 timepoint represents 20 minutes of GAL induction with no cyloheximide. **B)** Graphical representation of the fraction degraded, with data from **A)**.

**Fig. S9.**
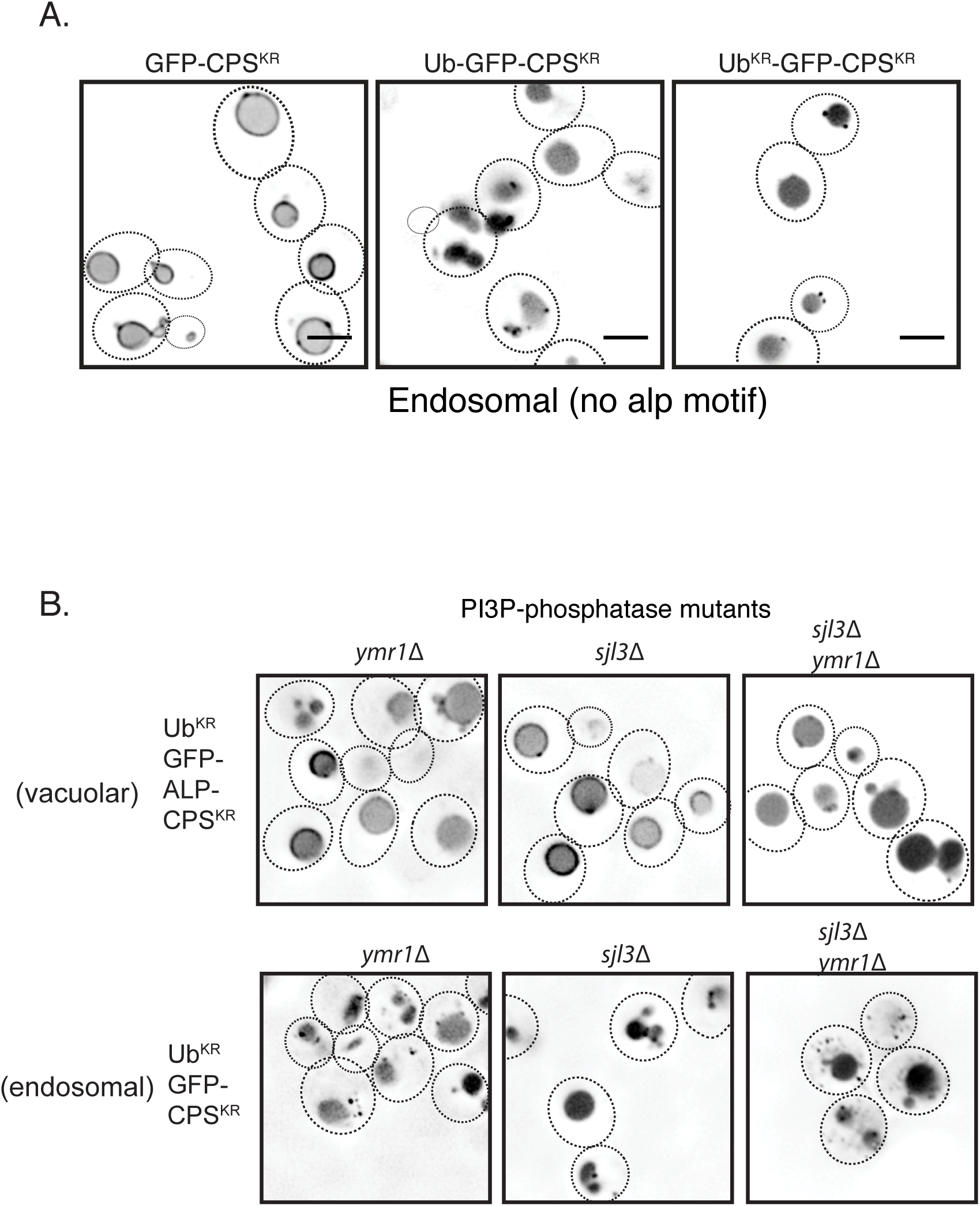
**A)** Localization of the GFP-CPS constructs with the K8R K12R (KR) mutations, when conjugated with ubiquitin or Ub^KR^. Ub^KR^ has all the lysines mutated to Arg. CPS^KR^ represents two mutations: K8R and K12R in the cytosolic region of CPS. B) Localization of the different CPS constructs in the phosphatase mutants *ymr1*Δ and *sjl3*Δ.

**Fig. S10.**
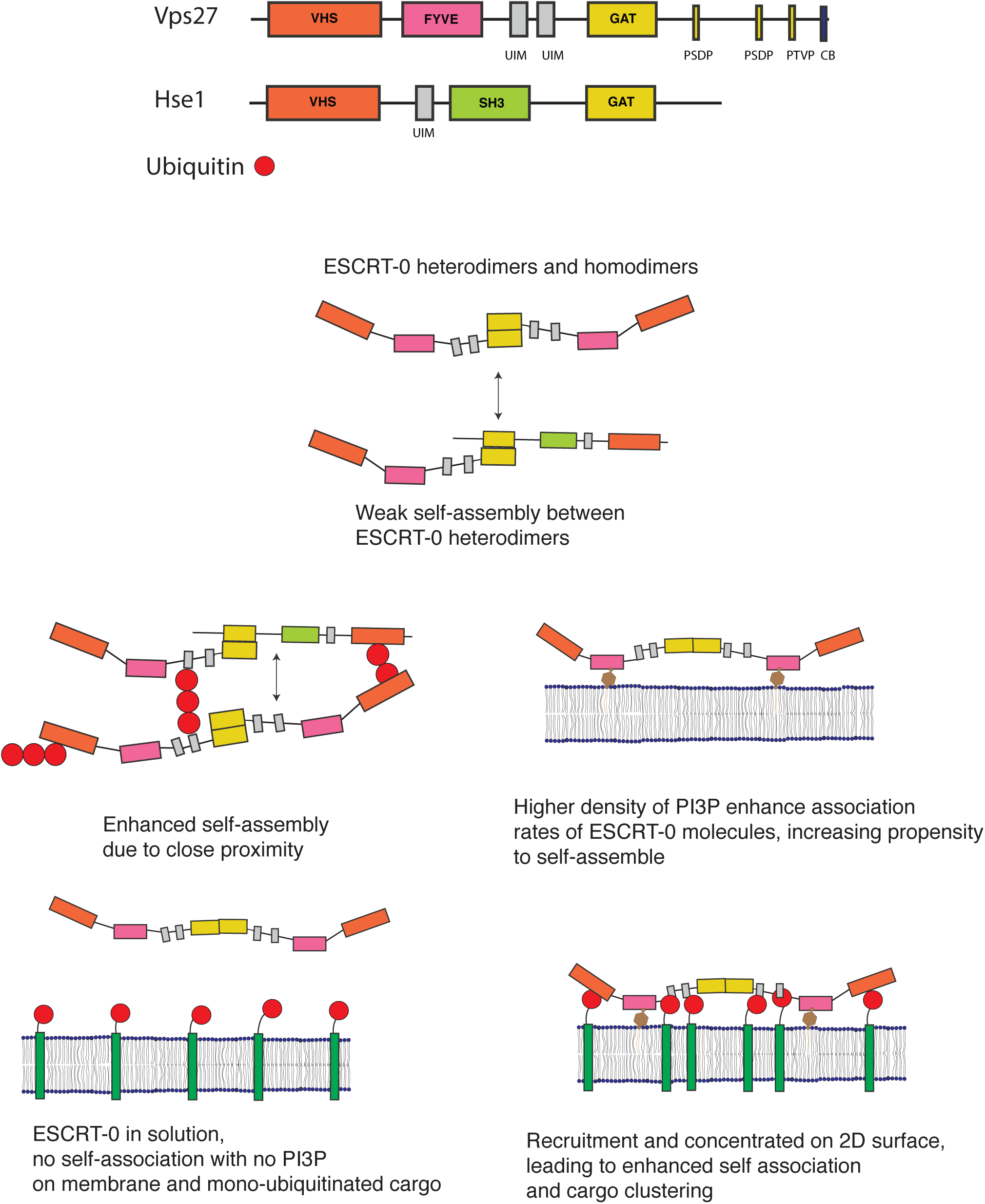
ESCRT-0 and ubiquitin assembly models on the surface of membranes. Surface density as regulated by lipid species, multivalency through polyubiquitin and multiple ubiquitin binding sites, and self-assembly through homo and hetero-oligomerization synergize to affect cargo condensation on membranes.

